# Small serine recombinases are markers for antiphage defense system discovery

**DOI:** 10.1101/2025.10.07.681051

**Authors:** Shelby E. Andersen, Joshua M. Kirsch, Navtej Singh, Stephen R. Garrett, John C. Whitney, Jay R. Hesselberth, Breck A. Duerkop

## Abstract

Renewed interest in phage therapy has highlighted a need to understand how bacteria subvert phage infection through antiphage defense systems. Traditionally, strategies to identify antiphage defense systems lack throughput or have limitations for bacterial species where antiphage defense systems are understudied. Herein, we developed a bioinformatic pipeline that uses a small serine recombinase to identify known and unknown antiphage defense systems. Using this approach to query reference genomes and metagenomes, we show that small serine recombinase genes are genetically linked to antiphage defense systems and serve as bait for finding these systems across diverse bacterial phyla. Using co-transcription predictions and statistical analysis of protein domain abundances, we experimentally validated our bioinformatic approach by discovering that KAP P-loop NTPases are fused to putative antiphage domains and reinforce prokaryotic Schlafen proteins as a new class of antiphage defense. Our work shows that small serine recombinases are a reliable genetic marker for the discovery of antiphage defenses across diverse bacterial phyla.

## Introduction

Multi-drug resistant (MDR) bacterial infections are a serious public health threat. With limited treatment options available for these infections, bacteriophage (phage) therapy has re-emerged as a potential treatment option^1–3^. Antiphage defense systems adapt to phages to overcome predation which threatens the efficacy of phage therapies^4^. Therefore, understanding the breadth, activity, and evolution of antiphage defenses in bacteria will inform the development of successful phage therapies.

Antiphage defense systems are routinely discovered by the tendency of these systems to localize in so-called ‘defense islands’, wherein uncharacterized phage defense systems are identified by their proximity to other known defense systems, or by generating genomic libraires and screening these in a heterologous bacterial host for phage defense^5–9^. To date, a marker gene that is broadly associated with various antiphage defenses across diverse bacteria has not been identified. Additionally, our knowledge of antiphage defense systems stems mostly from model bacteria such as *Escherichia coli* and *Bacillus subtilis*^5,8,10–13^. Well-studied antiphage defenses, such as abortive infection, restriction modification, and CRISPR/Cas, are widespread throughout bacterial phyla; however, the majority of the more recently discovered antiphage defense systems are primarily identified in the Pseudomonadota (formerly known as Proteobacteria) and sometimes poorly represented among diverse phyla^14^. This indicates that utilizing defense islands or random cloning as the primary means of discovering novel systems likely limits the discovery of antiphage defense biology and necessitates deeper studies across the bacterial domain of life.

Mobile genetic elements (MGEs), including plasmids, temperate phages, and transposons are significant drivers of genetic diversity in bacteria. MGEs are capable of spreading genetic traits that support bacterial fitness, such as antibiotic resistance genes and bacterial immunity genes to toxins^15,16^. Recent studies show that antiphage defense systems localize within the boundaries of MGEs^17–20^. For example, temperate phages and other phage-like elements are “hotspots” for antiphage defenses, and mobilizable antibiotic resistance cassettes can harbor and disseminate antiphage defenses^10,21^. Finally, genes that support DNA mobilization including recombinases, integrases, and transposon-encoding insertion sequence elements are often embedded within or near antiphage defense islands^19,22^. This indicates that broadly conserved MGE-specific genes could be leveraged as bait to identify antiphage defenses beyond those already discovered.

We recently characterized a Type IV restriction enzyme (TIV-RE, EF_B0059) from *Enterococcus faecalis* that is encoded on a bicistronic mRNA with a small serine recombinase (EF_B0058)^23^. The TIV-RE restricts phage replication in *E. faecalis*, while the co-transcribed serine recombinase is dispensable for phage resistance, suggesting that instead, the serine recombinase has evolved to perform an unknown function associated with the genetic maintenance of this TIV-RE defense system. Thus, we hypothesized that other antiphage defense systems could be found in the genomic regions surrounding homologs of these small serine recombinases and may also be co-transcribed. To explore this idea, we developed a bioinformatic pipeline termed Recombinase Associated Defense Search (RADS) that searches for small serine recombinase homologs and captures their surrounding genomic context. Using the RADS pipeline to query bacterial genomes and metagenomic contigs, we identified a plethora of antiphage defense systems. RADS identifies antiphage defenses spanning diverse bacterial phyla. Using binomial statistical testing coupled with co-transcription prediction to prioritize genes to further interrogate, we discover previously uncharacterized antiphage defense systems. We show that KAP NTPase domains regulate the function of linked modules that mediate antiphage defense. Because it relies on a conserved small serine recombinase rather than known antiphage defense genes, RADS is a unique method for locating antiphage defense systems in diverse bacteria without relying on homology to known defense systems.

## Results

### Development of a bioinformatic pipeline that identifies antiphage defense systems

The observation that a TIV-RE defense system (EF_B0059) in *Enterococcus faecalis* is encoded on a shared mRNA transcript with a predicted small serine recombinase (EF_B0058, **Fig. 1A**)^23^ led us to hypothesize that serine recombinases may co-occur in genomic regions with antiphage defense genes. To test this, we devised a bioinformatic approach that identifies small serine recombinases homologous to the *E. faecalis* serine recombinase EF_B0058, extracts the surrounding DNA sequence, and predicts the surrounding open reading frames (ORFs) that are then assessed for antiphage defense genes (**Fig. 1B**). We call this bioinformatic strategy Recombinase Associated Defense Search (RADS).

**Figure 1.**
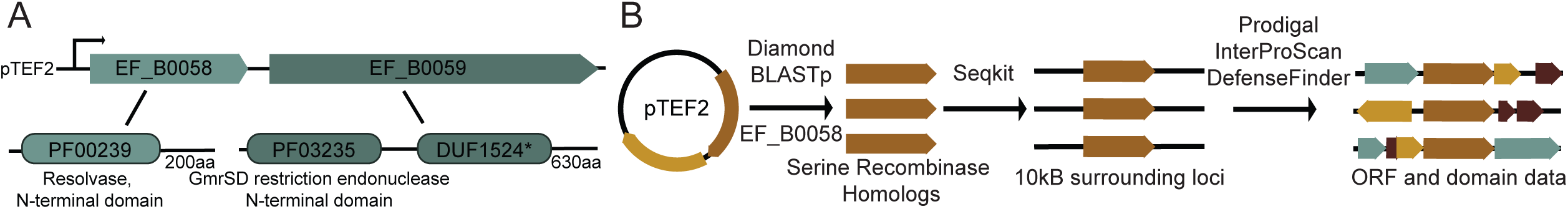
RADS leverages small serine recombinases associated with antiphage defense. (A) Graphic depicting the locus that inspired RADS. EF_B0058 is a serine recombinase that is co-transcribed with EF_B0059, a Type IV restriction enzyme on the large conjugative plasmid pTEF2. (B) Graphic depicting the RADS bioinformatic pipeline.

*E. faecalis* belongs to the Bacillota phylum. Therefore, to test whether RADS could identify previously discovered antiphage defense genes, we ran the pipeline on all complete genomes belonging to the Bacillota from Refseq (accessed March 2026). RefSeq was used because it ensures complete and unique genomes are being searched. RADS identified 9,478 EF_B0058-like serine recombinases in 13,553 genomes, resulting in a discovery rate of 0.215 hits per Mb (0.699 hits per genome). We next sought to provide evidence that the homologs of EF_B0058 identified by RADS were likely to be proteins with demonstrated small serine recombinase activity. Therefore, we took the nine lowest homology matches to EF_B0058 and ran Alphafold3 modelling on these homologs and EF_B0058^24^. These predicted structures were aligned to the predicted structure of the *Escherichia* phage Mu serine recombinase – a previously characterized small serine recombinase (**Fig. 2A**)^25^. The predicted structures of these proteins are nearly identical with an average RMSD value of 0.939Å, suggesting that these are similar small serine recombinases, despite their diverse sequences. Phylogenetic analysis reveals that the nine lowest-homology EF_B0058 homologs identified in the Bacillota dataset cluster with EF_B0058 and independently from large serine or tyrosine recombinases. (**Fig. 2B)**.

**Figure 2.**
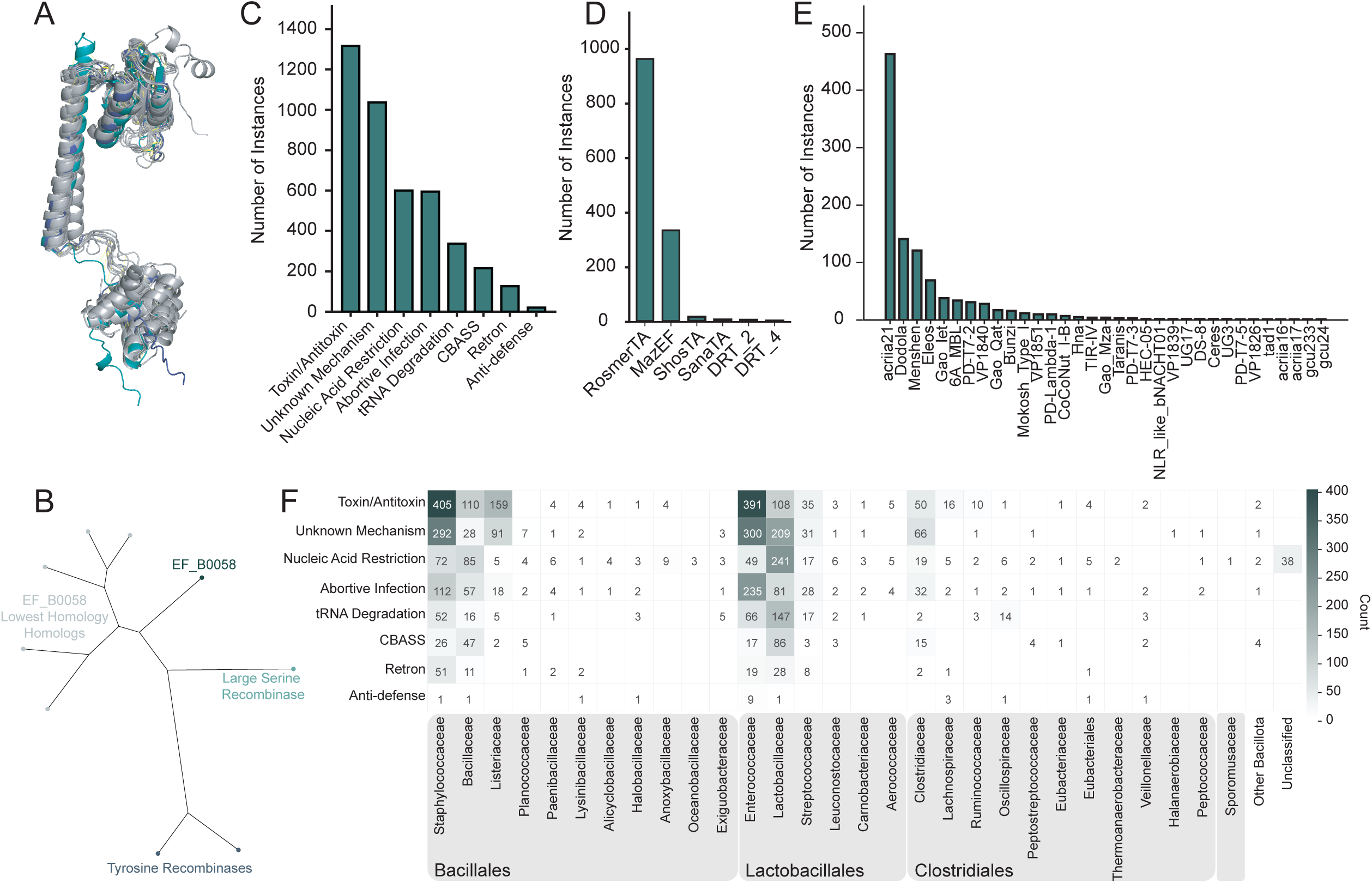
RADS identifies known antiphage defenses across genera in the Bacillota phylum. (A) Alphafold3 modeling of EF_B0058 (navy) and its 9 lowest-homology homologs from the Bacillota dataset (gray) aligned to the *Escherichia* phage Mu small serine recombinase (aqua); RMSD=0.939. (B) Unrooted phylogenetic tree of 9 lowest-homology EF_B0058 homologs identified in the Bacillota phylum. A large serine recombinase from *E. faecalis* and two representative tyrosine recombinases, one from Bacillota and one from Bacteroidota, are included for reference. Some low-homology homologs were identical in sequence, resulting in 5 homolog branches. (C) Quantification of antiphage defense system categories, (D) toxin/antitoxin types, and (E) unknown mechanism types identified by DefenseFinder. (F) Heatmap of antiphage defense system categories identified across families in the Bacillota phylum. Gray boxes indicate bacterial orders. The data supporting this figure can be found in S1 Data.

To assess whether the contigs identified by RADS (hereafter referred to as RADS contigs) had known antiphage defense systems we used DefenseFinder to query these DNA sequences^14^. Previously characterized antiphage defense systems were grouped into one of eight categories: abortive infection, CBASS, nucleic acid restriction, toxin/antitoxin, retron, tRNA degradation, anti-defense, and unknown mechanism. Known antiphage defense systems were found in high abundance within these contigs at a discovery rate of 0.681 hits per contig (**Fig. 2C**). The most abundant antiphage defense category identified among the Bacillota were toxin/antitoxin systems (**Fig. 2C)**, composed primarily of RosmerTA systems and others (**Fig. 2D**). The second most common category identified was unknown mechanism **(Fig. 2C)**, which exhibited a wide array of systems where knowledge of mechanism of action is limited or remains to be determined (**Fig. 2E)**.

Traditional antiphage defense system search methods have been historically limited in the diversity of bacteria interrogated. We sought to demonstrate that RADS could identify known antiphage defense systems in diverse bacteria. To first illustrate this, we assessed the distribution of known antiphage defense systems across the Bacillota phylum. Antiphage defense systems were observed in contigs from 29 bacterial families, spanning four orders (**Fig. 2F**). These data show that RADS can identify characterized antiphage systems in bacteria that have been previously neglected for antiphage defense discovery.

### RADS identifies an abundance of antiphage defense systems from microbiomes

To test the utility of RADS for finding antiphage defense systems in diverse phyla of bacteria from polymicrobial communities, we ran the pipeline on a large publicly available human fecal microbiome dataset^26^. We acquired 711 metagenome assemblies from pooled and un-pooled samples (encompassing a total of 2,441 samples) from the Human Microbiome Project Data Analysis and Coordination Center (HMP-DACC)^26^ (accessed July 2024). Using these data, we asked whether small serine recombinases were distributed evenly across phyla. We first determined the bacterial taxonomy using kraken2 with the phanta database^27^, revealing Bacillota (67.14%), Bacteroidota (24.23%), Pseudomonadota (3.21%), Uroviricota (2.26%), and Actinomycetota (1.75%) to be the most abundant phyla (**Fig. 3A**). Contigs containing moderate homology to the EF_B0058 serine recombinase (≥30% amino acid identity) were identified using RADS with a discovery rate of 0.016%. Serine recombinase-containing contigs shared a similar taxonomic distribution compared to the input dataset, revealing contigs from Bacillota (69.41%), Bacteroidota (21.34%), Pseudomonadota (3.26%), Uroviricota (3.03%), and Actinomycetota (2.27%) to be most abundant (**Fig. 3A**). This shows that small serine recombinases are well distributed across microbial phyla and that RADS can identify serine recombinase-associated loci across diverse bacteria.

**Figure 3.**
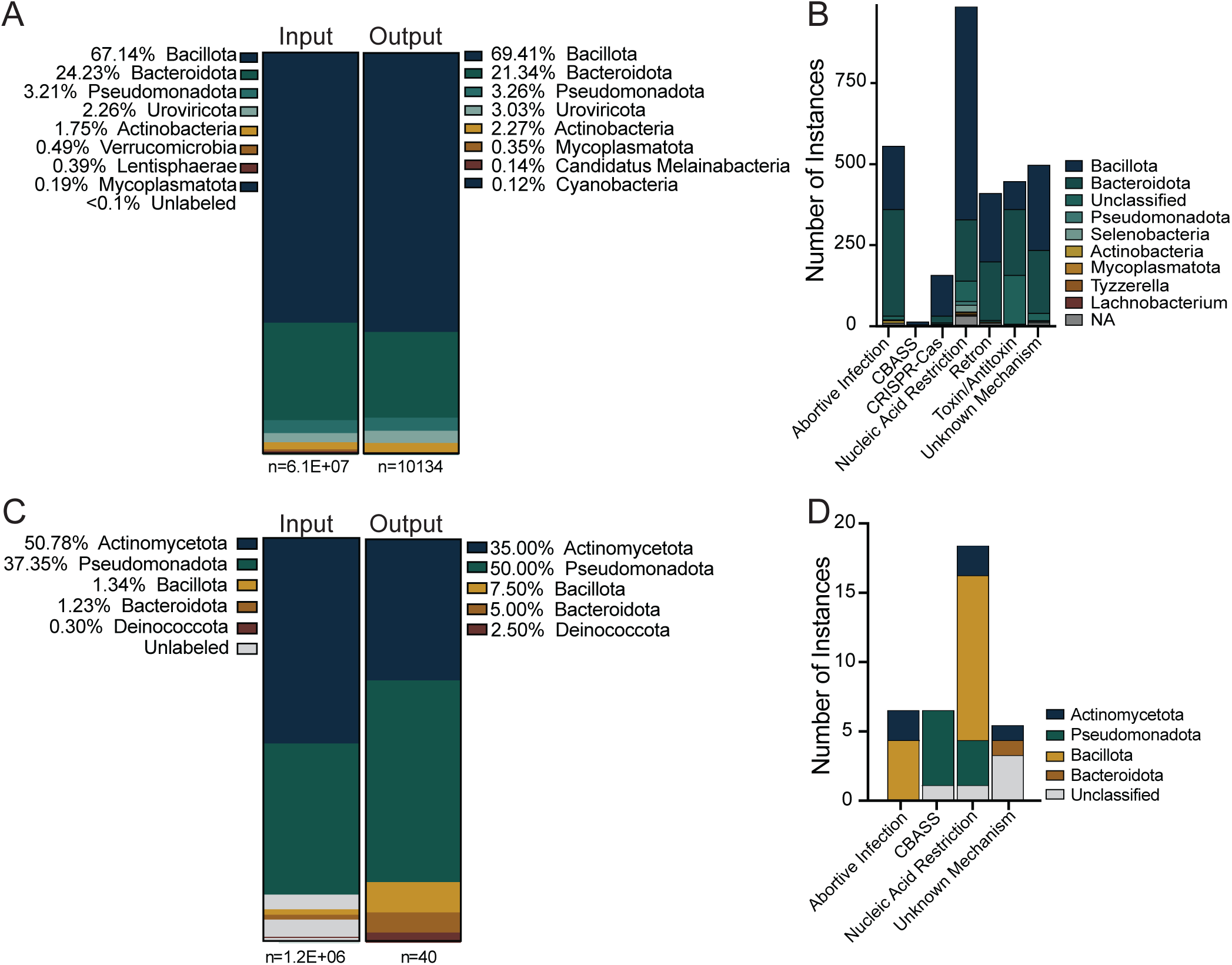
RADS identifies known antiphage defenses across diverse phyla. (A,B) Human fecal microbiome metagenomes or (C,D) soil microbiome metagenomes were run through the RADS pipeline. (A,C) Distribution of phyla identified by Kraken2 in the input dataset vs the contigs that came out of the RADS pipeline (output). (B,D) Quantification of antiphage system categories identified by DefenseFinder, where fill color represents the distribution of that category across different phyla. The data supporting this figure can be found in S1 Data.

We next sought to determine if RADS could identify known antiphage defense systems in the diverse phyla from these metagenomes. Abortive infection, CBASS, CRISPR-Cas, nucleic acid restriction, retron, toxin/antitoxin, and systems of unknown mechanism were identified by Defense Finder HMMs across nine phyla at high abundance with an antiphage discovery rate of 30.2% in RADS contigs (**Fig. 3B**). Notably, the distribution of phyla in which antiphage systems were identified is similar to the distribution of phyla in the input and output datasets.

To demonstrate if RADS is broadly applicable to bacteria from different environmental niches, we ran the pipeline on a previously published soil microbiome dataset composed of 3,304 metagenomes^28^. RADS contigs were identified at a discovery rate of 0.003%. While this discovery rate is approximately five-fold lower than observed for the gut microbiome dataset, this may be explained by contig quality of the input data. The average contig length for the soil metagenomes was 249.6bp, with a maximum length of 7,103bp. Meanwhile, the average and maximum length of the HMP-DACC dataset was 1796bp and 186,996bp, respectively. Importantly, input and output phyletic distributions were similar in the soil metagenomes with Bacillota being a minority in the output dataset (**Fig. 3C**). We identified 34 antiphage defense systems by Defense Finder HMM in these RADS contigs, with a discovery rate of 85% (**Fig. 3D**).

To further substantiate that RADS can identify antiphage defenses in phyla divergent from the Bacillota, we ran RADS on genomes from the Bacteroidota. We pulled all complete Bacteroidota genomes from NCBI Refseq (accessed March 2026). Of the 2,035 genomes searched, 278 RADS contigs were discovered, resulting in a discovery rate of 0.035 hits per Mb (average 0.143 hits per genome). Known antiphage defense systems were readily identified at a rate of 0.877 systems per RADS contig (**Fig. 4A**), and these systems were distributed across 13 families encompassing five orders (**Fig. 4B**). The most abundant defense system category of the Bacteroidota were toxin/antitoxin systems followed closely by abortive infection and nucleic acid defense systems (**Fig. 4C, 4D**). Altogether, these data validate RADS as a robust method for identifying putative antiphage defenses across highly diverse bacterial genomes.

**Figure 4.**
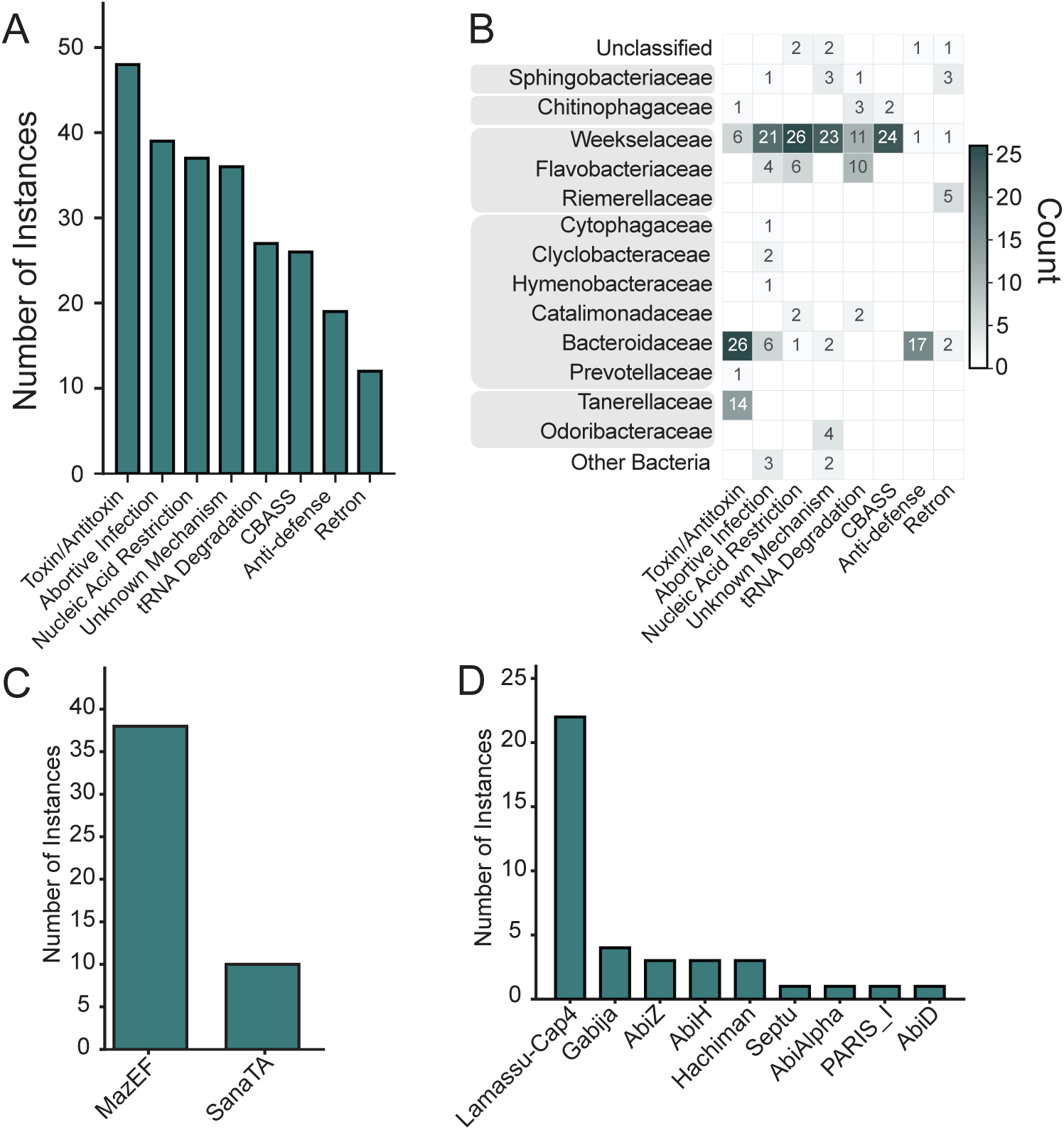
RADS identifies known antiphage defenses in the Bacteroidota. (A) Quantification of antiphage defense categories identified in the Bacteroidota. (B) Distribution of antiphage defense categories across families in the Bacteroidota. Gray boxes around family names indicate taxonomic orders. (C,D) Quantification of systems identified by DefenseFinder that are in the (C) toxin/antitoxin or (D) nucleic acid restriction categories. The data supporting this figure can be found in S1 Data.

With the success of using the EF_B0058 serine recombinase as bait for identifying antiphage defenses, we questioned whether this may be a general feature of recombinases. To test this, we utilized a tyrosine recombinase as bait. Similar to the serine recombinases, tyrosine recombinases are widely distributed in prokaryotic genomes and are involved in transposon mobility and prophage integration^29^. We ran RADS with either the *Bacteroides fragilis* tyrosine recombinase *tsr25*^30^ or a *Clostridia* tyrosine-type recombinase/integrase (NCBI accession MFQ9799422.1). This recombinase was selected from *Clostridia* because it was the most closely related tyrosine recombinase to *tsr25* in the Bacillota phylum. Using these tyrosine recombinases as bait, RADS was then run on either the Bacillota phylum or 33,520 randomly selected Bacteroidota genomes from Genbank (**Fig. S1)**. Notably, these tyrosine recombinases exhibited phylum bias with very few homologs of both baits found in the Bacillota and a multitude of homologs of both baits found in the Bacteroidota dataset. When homologs were identified, an overwhelming abundance of antiphage defenses were noted in their surrounding 10kb contig, particularly restriction enzymes (**Fig. S1**). The same antiphage defenses were identified by both baits in the Bacteroidota phylum. This data highlights that while recombinases more generally may be found surrounding antiphage defense systems, small serine recombinases exhibit unique phylum-level impartiality.

### Binomial and co-transcriptional analyses reveal novel antiphage defense genes using RADS

Knowing that we could bioinformatically identify previously characterized antiphage defense systems using RADS, we next sought to determine if RADS could be used to identify uncharacterized antiphage defense systems. We focused our efforts on the dataset generated from the Bacillota phylum (**Fig. 2**). To prioritize potential antiphage defense systems for further investigation, we employed a binomial statistical analysis paired with co-transcriptional prediction between the serine recombinase gene and genes containing one or more assigned pfam domain (identified by InterProScan, **Table S2**). The co-transcriptional analysis determines if an ORF following a serine recombinase gene is on the same DNA strand and if its predicted start site is within the broad distance of 100bp of the end of the serine recombinase^31,32^ **(Fig. S2A)**. If both conditions are met, the ORF is flagged as likely to be co-transcribed. The binomial statistical analysis reports the enrichment of a given domain within a RADS contig compared to the entire pangenome of the input dataset **(Fig. S2B)**. ORFs that are predicted to be co-transcribed with their associated small serine recombinase were assigned Defense Scores (see Materials and Methods) based on their proximity to known antiphage defenses as a metric for whether these ORFs would have been discovered in traditional guilt-by-association studies. To our surprise, the co-transcribed ORFs found in the Bacillota phylum have a median Defense Score of 0.0, indicating they are likely to be missed in traditional antiphage searches **(Fig. 5B)**. For all domains found in ORFs predicted to be co-transcribed with a small serine recombinase, we also calculated family-level defense score statistics (Fig. 5D). In short, all ORFs with a given domain within recombinase-containing Bacillota genomes were identified and used as an anchor for a 10kb contig (5kb on each side of the ORF). Those contigs were then queried for antiphage defense systems. Each anchor ORF with the domain of interest was then assigned a Defense Score. All ORFs with a specific domain were treated as a family to assign a minimum, maximum, and mean Defense Score for that family. Nearly all domains produced a low Defense Score, with the median mean family Defense Score being 0.01 **(Fig. 5D)**. Domains whose family had a mean >0.5 included those known to be involved in CBASS, abortive infection, and prokaryotic Argonaute systems. Notably, within-family Defense Scores had wide ranges, often ranging from 0.0 to 1.0 **(Fig. 5E)**. This highlights the utility of RADS, as it provides a criterion to identify putative antiphage defense systems within genomes without relying on common antiphage domains or presence in defense islands.

**Figure 5.**
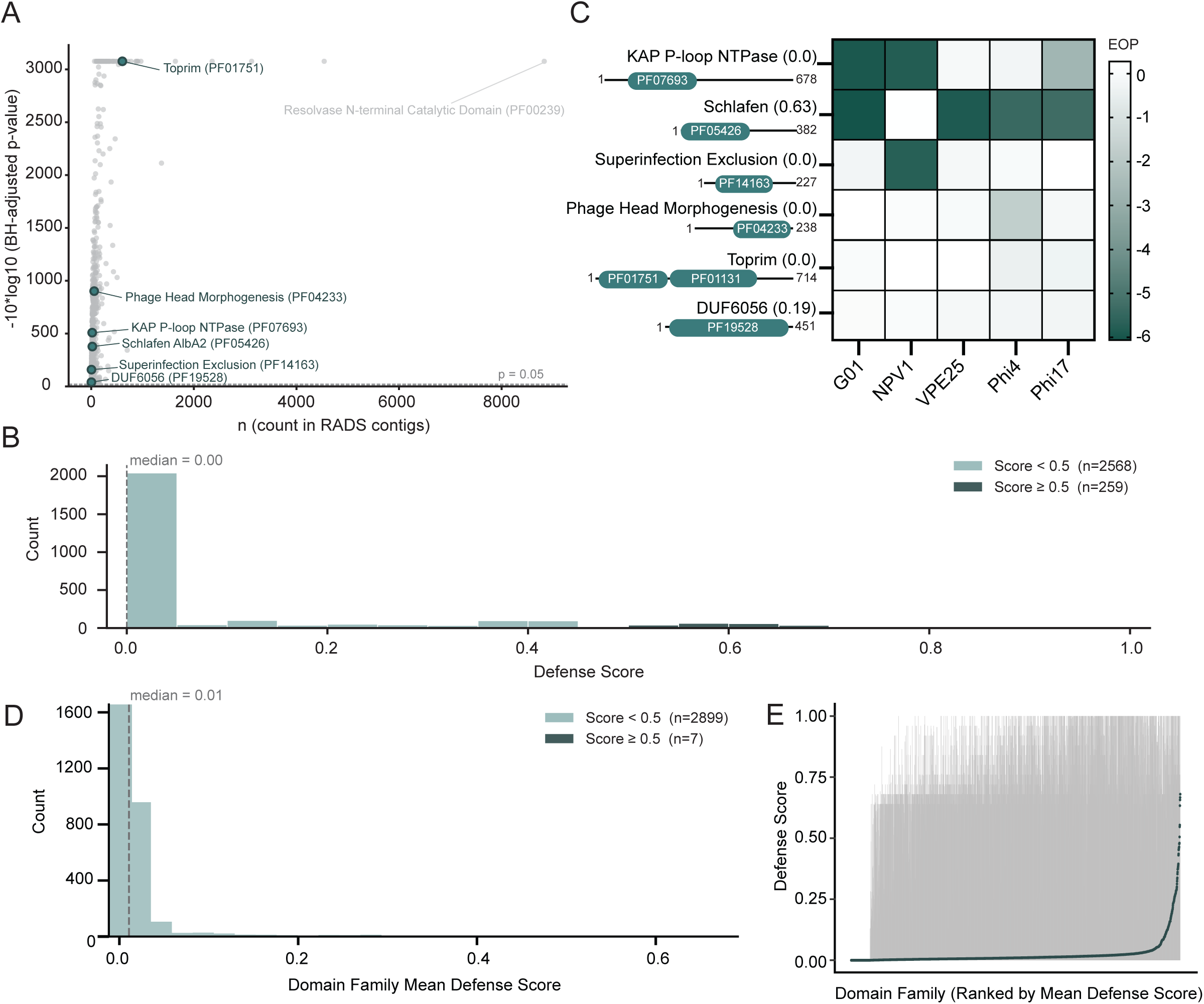
Discovery of previously uncharacterized antiphage defenses in RADS contigs. (A) Scatter plot of domains found in RADS contigs where n represents the number of instances in RADS contigs. The higher the value on the y axis, the more statistically enriched the domain. Green points represent domains chosen to test. (B) Histogram of defense scores of co-transcribed genes from the Bacillota dataset. A score below 0.5 indicates that the ORF is not likely to be in a defense island, whereas >0.5 indicates the ORF is likely in a defense island. (C) Heatmap of mean Log_10_ (fold-change efficiency of plaquing (EOP)) of phages (X-axis) on *E. faecalis* OG1RF carrying an ORF of interest (Y-axis) compared to strains carrying an empty vector. The more negative the EOP value, the stronger the antiphage defense. Defense Score for each transformed ORF is shown in parentheses. Visualization of the ORF transformed is displayed below each ORF label. Green regions represent domains with the Pfam accession listed within. (D) Histogram of mean domain family defense scores of genes containing a domain identified in an ORF co-transcribed with a small serine recombinase homolog (domains in panel 5A). A score below 0.5 indicates that the domain is not frequently found in defense islands, whereas >0.5 indicates the domain is often in a defense island. (E) Dot plot of mean domain family defense scores with gray lines representing the range of defense scores found in that family. Domain families are ranked by mean defense score on the X axis. The data supporting this figure can be found in S1 Data.

Six ORFs predicted to be co-transcribed with their associated small serine recombinase and encoding for significantly enriched domains by binomial statistical testing using Benjamini-Hochberg-corrected P-values (**Fig. 5A, Table S3**) were selected for testing for antiphage defense activity (**Fig. 5C**). Tested ORFs include a gene encoding a domain of unknown function (DUF) 6056, and genes predicted to encode putative phage head morphogenesis, superinfection exclusion, Toprim DNA topoisomerase, Schlafen, and KAP (Kidins220/ARMS and PifA) P-Loop NTPase domains (**Table S1, S3**). We prioritized the selection of ORFs from *Enterococcus* or other lactic acid bacteria to maximize the likelihood of successfully expressing these proteins in our model bacterium *E. faecalis* OG1RF, a strain that lacks prophages and other MGEs^33–35^. ORFs were selected from four *Enterococcus* species; *faecalis, faecium, lactis,* and *avium,* as well as *Lactobacillus helveticus* (**Table S1**).

We cloned these candidate antiphage defense genes and constitutively expressed them in *E. faecalis* OG1RF. We tested a panel of five genomically diverse *E. faecalis* phages for their ability to infect these strains^36–39^. Plaque forming units (PFU) per mL were measured and compared between *E. faecalis* cells expressing a potential antiphage defense system and cells carrying the empty vector to generate a fold change Efficiency of Plaquing (EOP) (**Fig. 5C, S3A**). This experiment revealed that three of the six ORFs tested were potent antiphage defenses with EOP being decreased by up to six log-fold change. These included the KAP P-Loop NTPase, Schlafen, and superinfection exclusion domain containing proteins (**Fig. 5C, S3A).** An additional two showed weak to negligible antiphage defense; Toprim and DUF6056 domain containing proteins (**Fig. 5C, S3A**).

### KAP P-loop NTPase domains are fused to antiphage domains

As KAP P-loop NTPases have not been robustly explored in association with antiphage defense, we focused our efforts on this system. To further probe the function of the *E. faecalis* KAP NTPase ORF (EF_KAP), we tested this system against various multiplicities of infection (MOIs) of phage. We first determined that EF_KAP does not function through abortive infection as it is equally protective at low (1) and high (10) MOIs **(Fig. S4).** AlphaFold3 modeling of EF_KAP suggests it consists of two transmembrane helices, one at the N-terminus and one within the KAP NTPase domain **(Fig. 6A)**. KAP P-loop NTPases have been previously reported to regulate the function of diverse proteins across the tree of life^40^. Notably, in prokaryotes these enzymes are hypothesized to regulate membrane-associated signaling complexes involved in restriction of foreign DNA^40^. A KAP P-loop NTPase domain is found in PifA, a protein encoded by the enterobacterial F plasmid that restricts phage T7, however it has been noted that phage restriction is likely not the primary function of this protein nor is the mechanism of action known^41^. In the case of EF_KAP, the transmembrane and KAP NTPase regions are followed by a 322 amino acid region with no known domains or predicted similarity to proteins of known function. This suggests that this KAP-NTPase is fused to an uncharacterized antiphage domain that may be activated by KAP-NTPase activity.

**Figure 6.**
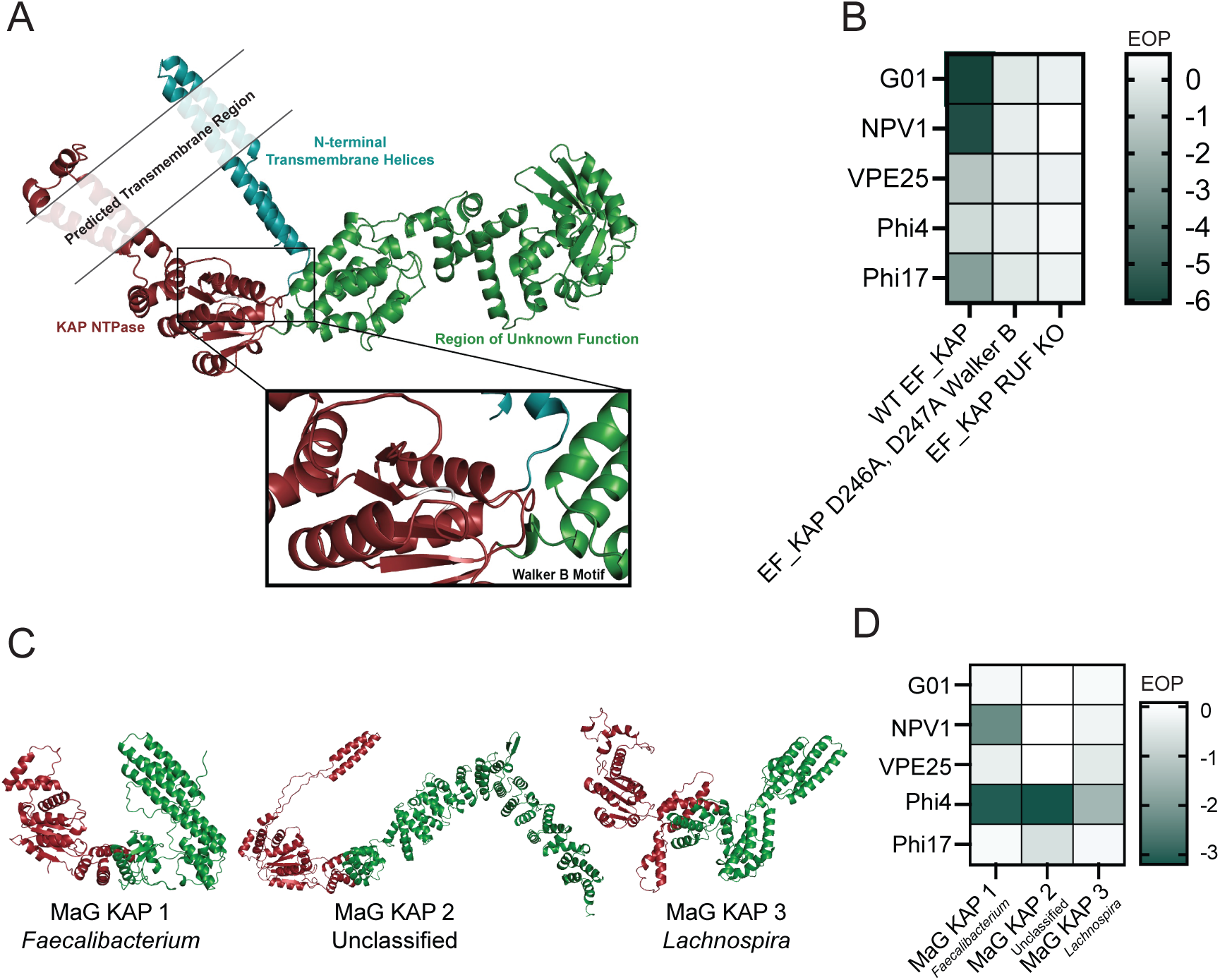
KAP P-Loop NTPases are fused to antiphage C-terminal regions. (A) Alphafold3 modeling of EF_KAP where aqua indicates the N-terminal transmembrane helices, red indicates the KAP NTPase domain, white indicates the Walker B motif within the KAP NTPase region, and green indicates the region of unknown function. (B) Heatmap of mean Log_10_ (fold-change efficiency of plaquing (EOP)) of phages (Y) on *E. faecalis* OG1RF carrying EF_KAP, EF_KAP with WalkerB mutations, or EF_KAP with the region of unknown function (RUF) deleted. The more negative the EOP value, the better the antiphage defense. (C) Alphafold3 modeling of KAP P-Loop NTPase domain (red) containing proteins with RUFs (green). (D) Heatmap of mean Log_10_ (fold-change efficiency of plaquing (EOP)) of phages on *E. faecalis* OG1RF expressing a different KAP P-Loop NTPase domain protein compared to strain carrying empty vector. The more negative the EOP value, the stronger the antiphage defense. The data supporting this figure can be found in S1 Data

To begin to unravel how EF_KAP mediates protection against phage infection, we generated two amino acid substitutions (D246A, D247A) in the Walker B motif of the KAP NTPase domain, which has been previously shown to be necessary for catalytic function by nucleotide hydrolysis^40^. Mutation of the Walker B motif ablated antiphage defense (**Fig. 6B, S3B**). These single amino acid mutations of the Walker B motif suggest that this protein is not preventing phage adsorption, but rather its enzymatic activity prevents phage replication. Deletion of the large region of unknown function also inactivates phage defense (**Fig. 6B, S3B**), demonstrating that both regions must work in concert to prevent phage infection.

We next tested whether KAP NTPase domains may regulate other fused regions for antiphage function and are divergent from EF_KAP. To explore this, we selected three KAP P-Loop NTPase domain-containing ORFs from the human fecal microbiome metagenomic RADS contigs with diverse C-terminal regions (**Fig. 6C**). These ORFs were selected from diverse host bacteria (*Faecalibacterium, Lachnospira*, and an unclassified contig) and had no predicted structural similarity to characterized proteins. Importantly, we found that these three ORFs conferred protection from phage infection in *E. faecalis* (**Fig. 6D, S3C**). Together, these data indicate that KAP P-Loop NTPases fused to regions of unknown function belong to an undescribed class of antiphage defense proteins.

## Discussion

In this study we developed a bioinformatic pipeline for identifying antiphage defense systems, which we have named Recombinase Associated Defense Search (RADS). This pipeline leverages a small serine recombinase as genomic ‘bait’ to identify adjacently encoded putative antiphage defense systems. RADS identifies antiphage defenses in diverse bacteria. We leveraged the predictive power of RADS to identify previously uncharacterized antiphage defense systems. Finally, our RADS bioinformatic pipeline provides query flexibility, thus the genomic region surrounding any gene of interest can be queried with minimal modifications.

Prior work has utilized bioinformatic strategies to identify antiphage defense systems^5–7^. However, these studies use known antiphage defense systems, prophages, or DNA genomic libraries to identify genomic regions that may assemble into defense islands^5–9,42^. Although an effective strategy, this approach is limited in the context of the genomic regions queried in a reference genome and is skewed toward specific bacterial species. Recently, two articles were published that leverage machine learning models to predict antiphage defense genes in bacterial genomes^43,44^. These models were trained on known antiphage defense systems identified by DefenseFinder and PadLoc^14,45^, and have further expanded our knowledge of both characterized and uncharacterized antiphage defense systems. Furthermore, Mordret *et al.*^44^ leveraged genomic proximity to known antiphage defenses and a large language model to deduce other genomic contexts, essentially expanding upon traditional guilt-by-association studies. RADS co-transcribed ORFs in the Bacillota exhibit extremely low Defense Scores, suggesting these systems would have been missed using such strategies, though it would be of interest whether small serine recombinases came up as a predictive marker in the latter study. Similar to the large language models recently developed, RADS represents an additional innovation for the identification of antiphage defenses to more evenly assess the presence of these systems in diverse bacterial phyla.

RADS discovers known antiphage defense systems at varying rates across the queried phyla – 0.681 hits per contig in the Bacillota compared to 0.877 hits per contig in the Bacteroidota. While this could point to an uneven association between small serine recombinases and antiphage defense systems between phyla, this could also be explained by the uneven discovery of antiphage defenses across phyla. Defense Finder is skewed toward identifying antiphage defense systems in the Pseudomonadota^14^, and homology-based searches are likely to perform less well the more disparate the bacterial species. This observation underscores the importance of discovery and functional elucidation of systems native to other phyla.

Nearly every category of antiphage defense system can be found in RADS contigs. However, toxin/antitoxin systems are the most commonly identified RADS-associated antiphage system in both the Bacillota and Bacteroidota. Toxin/antitoxin systems typically mediate abortive infection by disruption of transcription/translation of the antitoxin, allowing the toxin to kill the host cell^46^. Disruption of host gene expression is common in phage infection, as phages overtake host cell machinery to express their own genes. Because this gene expression disruption is a common signal during infection by diverse phages, toxin/antitoxin systems can be highly promiscuous in the phages they restrict^46^. Systems that are functionally promiscuous are more likely to be adaptable if mobilized to a new bacterial host. Interestingly, CRISPR-Cas systems are not found in the Bacillota or Bacteroidota RADS contigs, and rarely in metagenomic contigs. CRISPR-Cas would not make a good mobilizable antiphage defense system, as they are highly specific to the phage targeted and require a large CRISPR array in order to function^47^. We postulate that small serine recombinases may function to facilitate the mobilization of antiphage defense genes and that more promiscuous antiphage systems would be favored for a fitness advantage to recipient cells.

RADS includes a ranking scheme for ORFs likely to be involved in antiphage defense through their statistical enrichment in contigs and their likelihood of being co-transcribed with a small serine recombinase gene based on genomic proximity. RADS streamlines the identification of candidate antiphage defense systems in a high throughput manner. Herein, we test six ORFs identified using RADS for their ability to protect against phage infection, discovering three highly potent and two moderately potent antiphage defense systems. However, this only begins to scratch the surface of ORFs present in these RADS contigs. There are 2083 domains that are enriched (adjusted p-value < 0.05) in Bacillota contigs that have potential to be antiphage defense systems, regardless of co-transcription status. Multiple ORFs with diverse sequences can contain identical domains, thus studying how a given domain functions across diverse ORFs could provide new insights into domain function. Finally, ORFs identified with minimal antiphage defense activity may simply not be active against the phages tested, or *E. faecalis* OG1RF is an unsuitable heterologous host.

RADS offers a path toward the continued discovery of new biology related to antiphage defense systems. While KAP P-loop NTPases were hypothesized to be involved in antiphage defense two decades ago^40^, no follow up work has been done to support this. Here we show proteins harboring the KAP P-loop NTPase domain are potent antiphage defenses. We discovered a variety of KAP P-loop NTPases that offer cross-genus protection against phages. These KAP P-loop NTPases are associated with regions that lack known domains or any structural similarity to characterized proteins. We also show that the KAP P-Loop NTPases described in our study likely constitute a new class of proteins where the KAP P-Loop NTPase domain is fused to an antiphage defense region. The regions of unknown function fused to these KAP P-loop NTPases are diverse and exhibit unique phage defense profiles, suggesting they may represent diverse antiphage mechanisms. Broadly, NTPase domains are now being studied for their association with antiphage domains. The protein Tmn, a YobI-like P-loop NTPase, with transmembrane helices following the Walker A motif, synergizes with the Gabija defense system^6,40,48^. Recent studies show that the Tmn system restricts phage replication by hydrolyzing ATP and causing plasmolysis^49^. Furthermore, NTPases are also associated with nucleases with promiscuous antiphage activity^50^. The observation that NTPases are associated with diverse antiphage systems, highlights the importance of conserving these domains to control phage defenses in a variety of bacteria.

Expanding on the discovery of previously unrecognized antiphage defense systems, we show that a Schlafen domain-containing protein from *E. lactis* is a broad and potent antiphage defense system. Schlafens are mammalian antiviral proteins with extensive activity against eukaryotic viruses^51^ (**Fig. S5A**). However, a recent study also identified Schlafen proteins in prokaryotes that support antiphage defense, independently corroborating our findings^52^. One such system mediates abortive infection via host bacterial tRNA cleavage upon recognizing a phage tail assembly protein^52^. This mechanism is likely conserved with eukaryotic Schlafen protein function where Schlafen domains are fused to sensor domains that trigger Schlafen antiviral activity (**Fig. S5A**). To date, characterized prokaryotic Schlafen proteins are fused to sensor domains that impart antiphage function (**Fig. S5A**). Intriguingly, the Schlafen protein identified in our study exhibits broad antiphage activity via abortive infection despite its C-terminus lacking any known domains **(Fig. S5A, S5B**). This suggests that a variety of yet to be described phage sensing mechanisms are mediated by Schlafen proteins, reminiscent of what has been observed in eukaryotes^33,51^.

Other antiphage defense systems identified in this study had phage-specific and/or moderate to weak effects on phage infection. This includes a predicted superinfection exclusion system and a phage head morphogenesis protein. Superinfection exclusion proteins support inter-phage defense, often encoded in prophages, that restrict invading phages^53^. Interestingly, the superinfection exclusion protein identified in our study is not found within a prophage as determined by PHASTER^54^. This may represent the hijacking of a superinfection exclusion mechanism without maintaining the prophage or potential recombination between a phage carrying this superinfection exclusion gene and the host bacterial chromosome. Further study of this superinfection exclusion mechanism may reveal unrecognized adaptive relationships between infecting phages and their host bacteria that promotes antiphage defense.

Due to the breadth of antiphage defense genes found associated with small serine recombinase genes, it is likely that this co-occurrence represents more than just evolutionary synteny. Rather, we propose that this link between small serine recombinase genes and antiphage defense systems represents a broader evolutionary adaptation. One hypothesis is that antiphage defense systems are mobilized by horizontal gene transfer via prophages or plasmids^17–20^. However, maintenance of such genetic elements can impose fitness constraints, such as the observation that plasmid maintenance can decrease bacterial growth rate^55^. We postulate that these small serine recombinases may help mobilize antiphage defenses to the chromosome, thereby minimizing the need for carriage of costly mobile genetic elements and reducing fitness costs. Additionally, small serine recombinases similar to the recombinase used in RADS, function as invertases that support DNA inversion events^56^. DNA inversion may be a means to regulate the expression of antiphage defense genes during phage infection. Notably, a recent study showed that the serine recombinase PinQ of the cryptic prophage Qin inverts DNA to form a protein coding sequence that supports antiphage defense by interfering with phage adsorption^57^. Finally, these small serine recombinases may serve to diversify the host genome upon phage predation while working in tandem with the antiphage defense system to block phage infection. Diverse populations are more resilient to infection^58^, and this could be a mechanism whereby genetic flexibility of phage defense systems through rapid recombination and horizontal gene transfer supports population resilience.

## Materials & Methods

### Plasmids

Plasmid constructs were generated by Genscript through synthesis of the nucleotide sequence of interest and cloning each gene into the plasmid pLZ12A harboring a constitutive promoter^39^. Site directed mutagenesis of antiphage defense genes were generated where mutations of interest were designed into the primers used for amplification. These primers were used for whole plasmid PCR amplification using Q5 High-Fidelity 2X DNA polymerase Master Mix (New England Biolabs) to create a linear DNA product. PCR products were purified using a QIAquick PCR Purification Kit (Qiagen) and linearized products were ligated using T4 DNA ligase (New England Biolabs). For two-piece assemblies, both products were digested with XhoI per the manufacturer’s recommendations (New England Biolabs) prior to ligation. Ligated plasmids were transformed into electrocompetent *E. coli* TG1 DUOs (Lucigen). Plasmids were purified from *E. coli* using a Plasmid Mini Prep Kit (Qiagen) and transformed into electrocompetent *E. faecalis* using the lysozyme treatment method^39^. Plasmids and primers used can be found in Supplemental Table 1.

### Bacteria and bacteriophages

A list of bacterial and bacteriophage strains used in this study can be found in Supplemental Table 1. *E. faecalis* strains were grown with aeration in Todd Hewitt broth (THB, Becton Dickinson) or on THB agar at 37°C. *Escherichia coli* was grown in Lennox lysogeny broth (LB, Fisher) with aeration or on LB agar at 37°C. When necessary, for the selection of *E. coli* or *E. faecalis* carrying a plasmid of interest, 15 µg/mL chloramphenicol (Research Products International) was added to the media. Phage plaque assays were performed using THB base and top agar supplemented with 10 mM MgSO_4_ as described elsewhere^38^.

### Bacterial growth curves

Overnight cultures of *E. faecalis* were grown in THB broth with 15µg/mL chloramphenicol. Overnight cultures were diluted to an OD_600_ of 0.3 in THB broth with 15µg/mL chloramphenicol and 10mM MgSO_4_. 200µL of the diluted culture was added to the wells of a 96-well plate. Phage diluted in SM-plus buffer (100 mM NaCl, 50 mM Tris-HCl, 8 mM MgSO_4_, 5 mM CaCl_2_ [pH 7.4]) was added to the appropriate samples. Samples were set up in technical triplicate. The 96-well plate was then moved to 37°C with orbital shaking. Bacterial growth was measured by optical density (OD_600_) every 5 minutes using a Biotek Synergy H1 Microplate reader.

### Bioinformatics

#### Development of the Recombinase Associated Defense Search (RADS) pipeline

Genomes of interest may be supplied to RADS or it has been outfitted with the ability to download sequences of interest from Refseq or Genbank. RADS first predicts ORFs in genomes or metagenomic contigs and translates them to amino acid sequences using prodigal^59^. Prodigal is used for de novo generation of ORFs within genomes and contigs rather than using the protein coding files from NCBI because the nascent ORF headers have the data in an easily queried format for downstream extraction. These amino acid sequences are then used to generate a BLASTp database using Diamond^60^. The Diamond BLASTp database is searched for homologs of the EF_B0058 small serine recombinase with moderate (≥30%) amino acid identity. Once the genomic coordinates of serine recombinase homologs are identified, Seqkit is used to extract the 5kB flanking DNA sequence up-and downstream of the serine recombinase gene (designated as a RADS contig)^61^. RADS can be tuned to extract a region of a preferred size, but defaults to 5 kb on each side of the serine recombinase gene for a total of 10kb. These contigs are then run through InterProScan, returning a table of putative domains and predicted structural similarities for each ORF across the 10kb region queried^62^. RADS will also generate a list of predicted co-transcribed ORFs in which the downstream ORF must be on the same DNA strand and within 100 base pairs of the serine recombinase gene (**Fig. S2A**). RADS has been packaged into a snakemake pipeline, allowing for easy configuration, process threading, and output data management. The configuration can be manipulated to control for all variables in the pipeline including the query used, dataset acquisition and usage, thresholds, and omission of computation-heavy metrics. Due to this packaging and configurability, RADS may be used to perform similar genomic analyses using other queries of interest with goals beyond the discovery of antiphage defense systems.

#### Analysis of RADS output data

The RADS pipeline has been outfitted to fully house the calculation and analysis of all output metrics. All metrics and output data can be viewed in the PyShiny interactive RADS dashboard. Notable features include: a key metrics page, a locus viewer with notable filters, domain exploration, co-transcription prediction, and the visualization of Defense Finder data.

The binomial analysis compares the frequency of a domain being found in a RADS contig to its frequency in the entirety of the genome to reveal domains that are truly enriched in RADS contigs (**Fig. S2B**). To run the binomial analysis, InterProScan must be run on full genomes where serine recombinase homologs were found. As this can be computationally expensive, this feature can be disabled via the configuration.

Known antiphage systems are confirmed using DefenseFinder^14^. Recombinase discovery rate was calculated by the number of recombinase homologs discovered divided by the number of input sequences. Recombinase discovery rate by megabase (Mb) was calculated by the number of recombinase homologs discovered divided by the total Mbs of input sequences. Antiphage discovery rate was calculated by the number of antiphage defense systems identified by DefenseFinder Genes divided by the number of contigs discovered by RADS. All protein modeling was performed with Alphafold3^24^.

For each ORF that is predicted to be co-transcribed with its associated small serine recombinase, RADS calculates a Defense Score. This Defense Score is a composite of a proximity score (0-1) and a density score (0-1). The proximity score is calculated by the distance in basepairs to the nearest Defense Finder gene on that contig. If the co-transcribed gene is found to be a defense system by Defense Finder, the proximity score is 1. As one gene is estimated to be 2000bp, if a defense system is found 2000bp away, the proximity score is set to 0.37. If no known antiphage defenses are found in that contig, the Defense Score is set to 0. The density score accounts for multiple antiphage genes on one contig. This metric is calculated as the count of the antiphage genes divided by 10 (density score = count / 10). The max genes for this metric is set to 10, meaning that 10+ defense genes found in the contig would result in max density. The proximity score and the density score are weighted slightly toward proximity (0.6X proximity, 0.4X density) and combined to net the composite score. A composite score of 0 suggests that this ORF would not have been found from traditional defense island searching, while a score >0.5 suggests that this ORF may be in a defense island.

For family-level defense scores, the InterPro domain IDs from the binomial analysis (IDs corresponding to Pfams found in ORFs predicted to be co-transcribed with their small serine recombinase) were used to extract all other ORFs with that domain found in the input whole genomes. ORFs sharing the same InterPro ID represent a protein family. Of note, only genomes with an EF_B0058 small serine recombinase homolog were used to pull these additional domain hits, as running InterProScan on all genomes in the phylum proved too computationally demanding. For each InterPro ID, a list of ORFs across all genomes was generated and 5kb on both sides of those ORFs were extracted. DefenseFinder was then run on these contigs and Defense Scores were calculated as indicated above. For each protein family, the minimum, maximum, and mean Defense Scores were calculated and reported.

#### Computational recommendations

As with all bioinformatic approaches, RADS exhibits computational limits. Paralellization via snakemake has made the analysis of large datasets timelier. The predicted relationship between number of genomes analyzed, available CPU, and time to completion can be found at the RADS Github or Zenodo repositories (see data availability section). Snakemake will automatically handle threading of tasks, and CPU for individual steps can be assigned during configuration.

#### Metagenomics analyses

Assembled metagenomes were downloaded from their respective data sources (Human Microbiome Project or Ma *et al*. 2023)^26,28^. Taxonomy classification was performed using Kraken2^27^. The human fecal microbiome metagenomes were taxonomically classified with the Phanta database, while the soil microbiome metagenome was taxonomically classified with the with the Plus_PF_16 database. These analyses were completed with the first version of RADS, prior to snakemake packaging. Antiphage defense systems were identified by Defense Finder HMM.

#### Phylogenetics

Amino acid sequences were aligned using mafft^63^. The phylogenetic tree was built by neighbor joining and optimized with maximum likelihood by Phangorn^64^. The tree was visualized with ggtree.

## Data Availability

The RADS pipeline is is available for download at https://github.com/Seandersen/RADS and can also be accessed at https://zenodo.org/records/21132456. RADS Bacillota and Bacteroidota datasets can be found at https://zenodo.org/records/21134694 and https://zenodo.org/records/21134503 respectively. The raw data associated with this study can be found in S1 Data.

---

**Figure S1. Tyrosine recombinases do not reliably identify antiphage defenses across phyla.** Tyrosine recombinases were used as queries in the RADS pipeline on the Bacillota and Bacteroidota phyla. Graphic depicts searches performed with number of hits (recombinase homologs) and antiphage systems discovered noted. Graphs represent the antiphage defense system categories identified in tyrosine recombinase contigs. The data supporting this figure can be found in S1 Data.

**Figure S2. Additional analyses utilized to select ORFs to test for antiphage activity.** (A) Graphic depicting the co-transcriptional analysis requirement for binomial analysis testing. ORFs were required to be on the same strand of DNA and within 100bp of the small serine recombinase to be flagged as likely to be co-transcribed with a small serine recombinase. (B) Graphic depicting the binomial analysis testing. The frequency of finding a given domain in RADS contigs versus the genome was compared to the overall distribution of domains in contigs compared to domains in the genome. Comparing these frequencies allows the assignment of a statistical metric for how enriched domains were in contigs.

**Figure S3. Quantitative bar plots of EOP experiments.** (A, B, C) Triplicate values (points) and means (bars) of EOP experiments in Figure 5B (A), Figure 6B (B), and Figure 6D (C). The data underlying this Figure can be found in S1 Data.

**Figure S4. EF_KAP does not protect from phage infection by abortive infection.** OD_600_ of *E. faecalis* OG1RF cultures was measured over time in the presence or absence of phage infection (phages G01, NPV1, and Phi17) at MOI of 1 or 10. EV indicates the *E. faecalis* carries empty vector, while EF_KAP indicates host strain carries plasmid expressing EF_KAP. Lines represent mean of two biological replicates of technical triplicates. The data underlying this Figure can be found in S1 Data.

**Figure S5. Schlafen proteins are conserved antivirals.** (A) Comparison between prokaryotic and eukaryotic Schlafen proteins of different groups. Asterisk indicates size of protein is not currently reported. (B) OD_600_ of *E. faecalis* OG1RF cultures were measured over time in the presence or absence of phage G01 infection at MOI of 0.1, 1, or 10. EV indicates host strain carries empty vector, while Schlafen indicates host strain carries plasmid expressing the Schlafen protein from this study. Lines represent mean of three biological replicates of technical triplicates

**Table S1. Bacterial/bacteriophage strains, plasmids, and oligonucleotides used in this study.**

**Table S2. Predicted co-transcription hits between small serine recombinase genes and genes containing one or more assigned pfam domain.**

**Table S3. Bacillota protein domains tested in for antiphage defense activity.**

