## Supplementary figures and images for "Small serine recombinases are markers for antiphage defense system discovery"

### Supplementary Figure 1

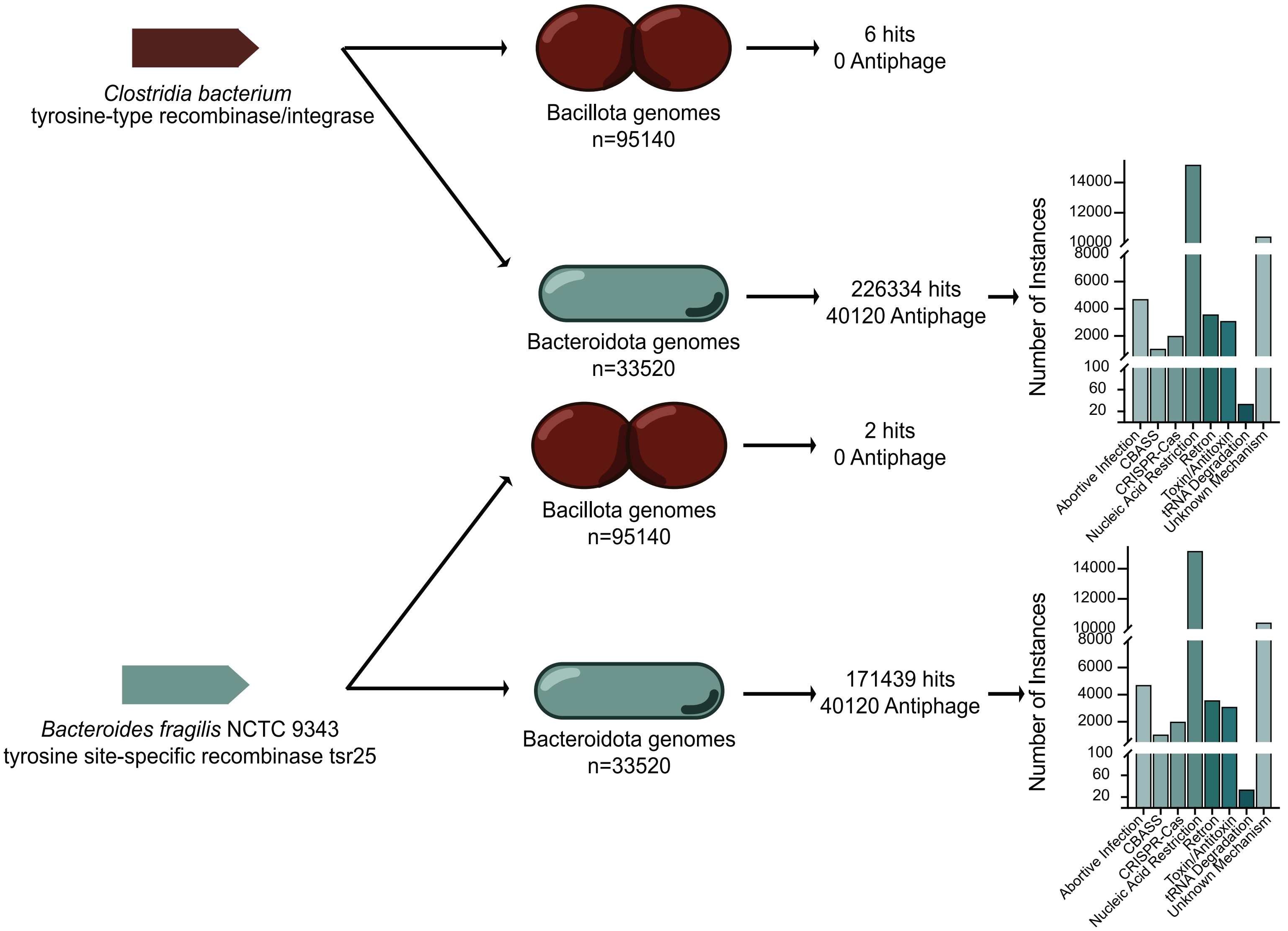

### Supplementary Figure 2

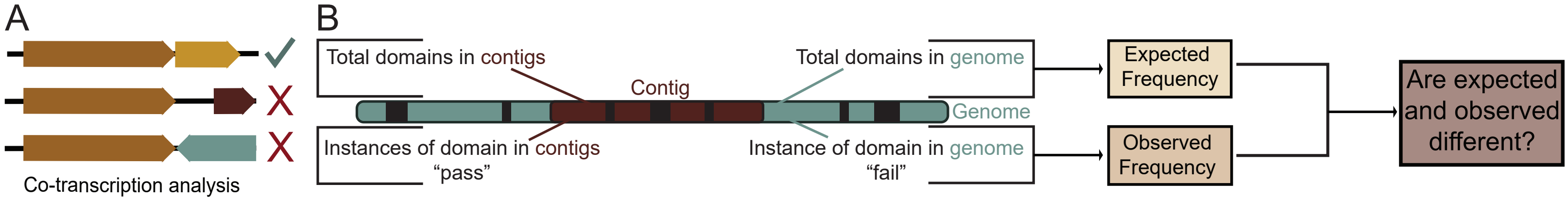

### Supplementary Figure 3

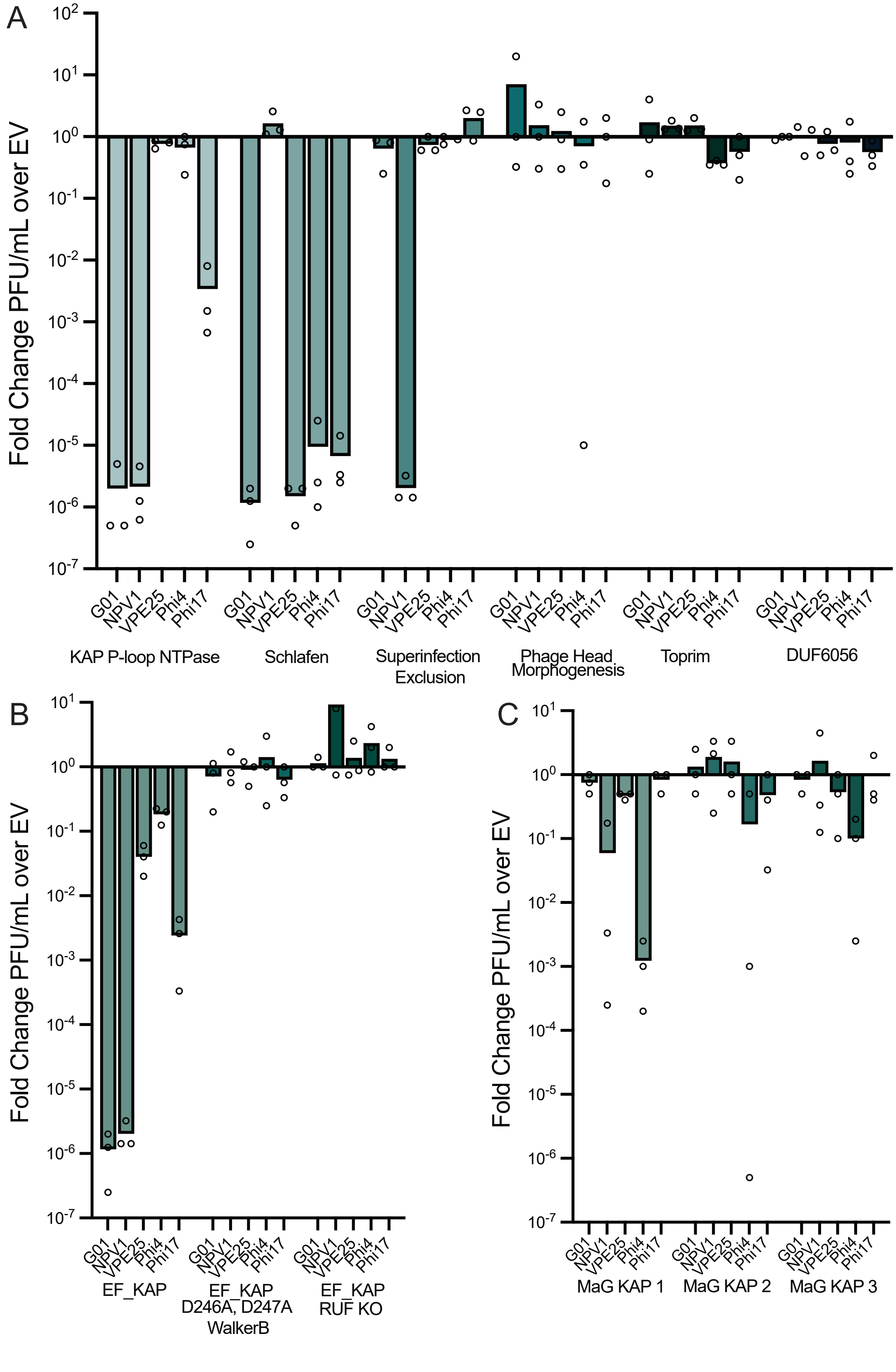

### Supplementary Figure 4

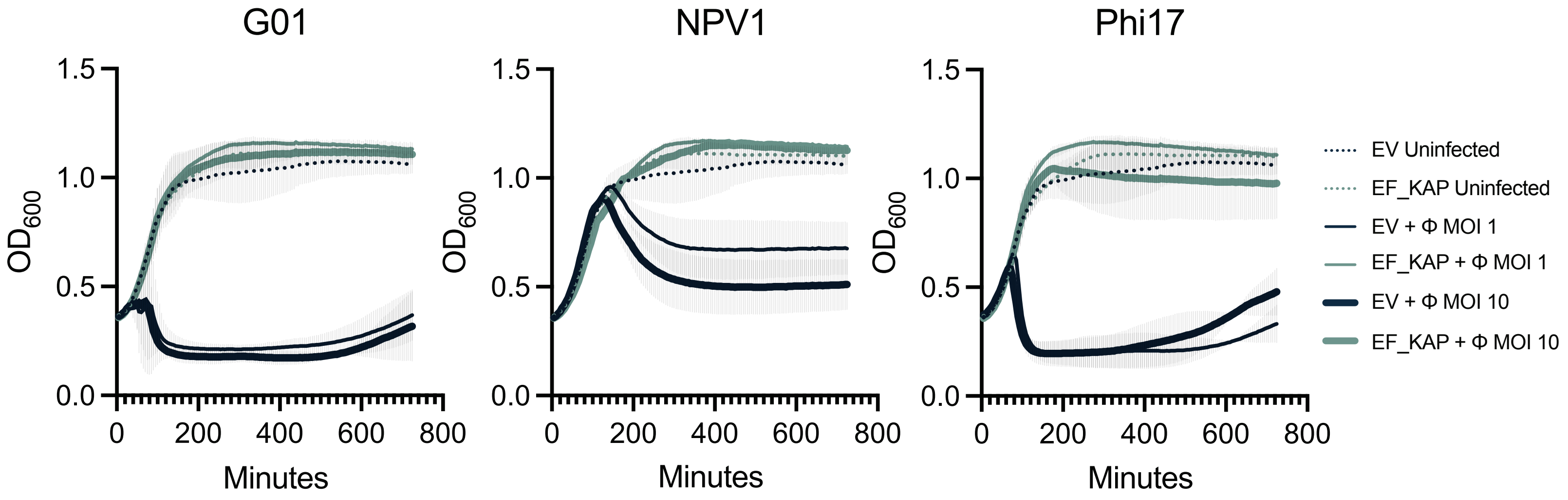

### Supplementary Figure 5

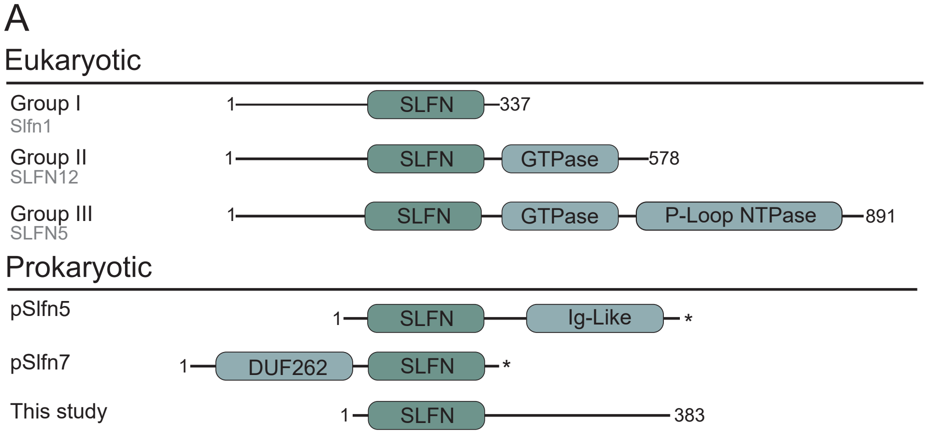

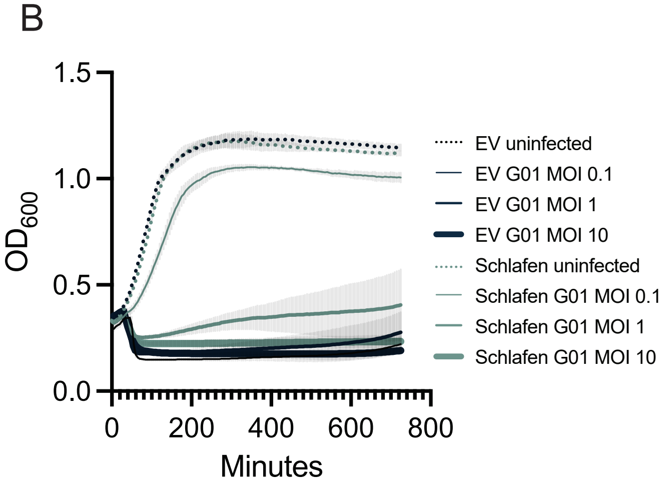
