## Supplementary Table 1 for "Small serine recombinases are markers for antiphage defense system discovery"

**Supplemental Table 1** Bacterial/bacteriophage strains, plasmids, and oligonucleotides used in this study.

| Category | Relevant Properties | Reference |
|---|---|---|
| **Plasmids** | | |
| pLZ12A | P-*bacA* promoter cloned into shuttle vector pLZ12; pSH71 origin; $Cm^R$ | Chatterjee & Johnson *et al.,* 2019 |
| pLZ12A::KAP NTPase / pLZ12A::EF_KAP | pLZ12A constitutively expressing EF_KAP; $Cm^R$ | This Study |
| pLZ12A::Schlafen | pLZ12A constitutively expressing Schlafen ORF; $Cm^R$ | This Study |
| pLZ12A::SuperInfection Exclusion | pLZ12A constitutively expressing SuperInfection Exclusion ORF; $Cm^R$ | This Study |
| pLZ12A::AAA ATPase | pLZ12A constitutively expressing AAA ATPase ORF; $Cm^R$ | This Study |
| pLZ12A::Phage Head Morphogenesis | pLZ12A constitutively expressing Phage Head Morphogenesis ORF; $Cm^R$ | This Study |
| pLZ12A::Abi | pLZ12A constitutively expressing Abi ORF; $Cm^R$ | This Study |
| pLZ12A::Toprim | pLZ12A constitutively expressing Toprim ORF; $Cm^R$ | This Study |
| pLZ12A::DUF5677 | pLZ12A constitutively expressing DUF5677 ORF; $Cm^R$ | This Study |
| pLZ12A::DUF6056 | pLZ12A constitutively expressing DUF6056 ORF; $Cm^R$ | This Study |
| pLZ12A::EF_KAP D246A,D247A | pLZ12A constitutively expressing EF_KAP D246A,D247A mutant; $Cm^R$ | This Study |
| pLZ12A::EF_KAP Effector KO | pLZ12A constitutively expressing EF_KAP Effector KO mutant; $Cm^R$ | This Study |
| pLZ12A::MaG KAP 1 | pLZ12A constitutively expressing MaG KAP 1 ORF; $Cm^R$ | This Study |
| pLZ12A::MaG KAP 2 | pLZ12A constitutively expressing MaG KAP 2 ORF; $Cm^R$ | This Study |
| pLZ12A::MaG KAP 3 | pLZ12A constitutively expressing MaG KAP 3 ORF; $Cm^R$ | This Study |
| **Strains** | | |
| *E. faecalis* | | |
| OG1RF | Human oral isolate; $Rf^R$, $Fa^R$ | Bourgogne *et al.,* 2008 |
| OG1RF pLZ12A | OG1RF carrying pLZ12A empty vector | This Study |
| OG1RF pLZ12A::EF_KAP | OG1RF carrying pLZ12A::EF_KAP | This Study |
| OG1RF pLZ12A::Schlafen | OG1RF carrying pLZ12A::Schlafen | This Study |
| OG1RF pLZ12A::SuperInfection Exclusion | OG1RF carrying pLZ12A::SuperInfection Exclusion | This Study |
| OG1RF pLZ12A::AAA ATPase | OG1RF carrying pLZ12A::AAA ATPase | This Study |
| OG1RF pLZ12A::Phage Head Morphogenesis | OG1RF carrying pLZ12A::Phage Head Morphogenesis | This Study |
| OG1RF pLZ12A::Abi | OG1RF carrying pLZ12A::Abi | This Study |
| OG1RF pLZ12A::Toprim | OG1RF carrying pLZ12A::Toprim | This Study |
| OG1RF pLZ12A::DUF5677 | OG1RF carrying pLZ12A::DUF5677 | This Study |
| OG1RF pLZ12A::DUF6056 | OG1RF carrying pLZ12A::DUF6056 | This Study |
| OG1RF pLZ12A::EF_KAP D246A,D247A | OG1RF carrying pLZ12A::EF_KAP D246A,D247A | This Study |
| OG1RF pLZ12A::EF_KAP Effector KO | OG1RF carrying pLZ12A::EF_KAP Effector KO | This Study |
| OG1RF pLZ12A::MaG KAP 1 | OG1RF carrying pLZ12A::MaG KAP 1 | This Study |
| OG1RF pLZ12A::MaG KAP 2 | OG1RF carrying pLZ12A::MaG KAP 2 | This Study |
| OG1RF pLZ12A::MaG KAP 3 | OG1RF carrying pLZ12A::MaG KAP 3 | This Study |
| *Phages* | | |
| G01 | *Enterococcus* phage isolated from Ganges River | Sheriff *et al.,* 2024 |
| Npv1 | *Enterococcus* phage isolated from sewage | Trotter & Dunny, 1990 |
| VPE25 | *Enterococcus* phage isolated from sewage | Duerkop *et al.,* 2016 |
| Phi4 | *Enterococcus* phage isolated by the Naval Medical Research Center (NMRC) | Chatterjee & Johnson *et al.,* 2019 |
| Phi17 | *Enterococcus* phage isolated by the Naval Medical Research Center (NMRC) | Chatterjee & Johnson *et al.,* 2019 |
| **Sequences** | | |
| *ORFs Tested* | | |
| KAP NTPase / EF_KAP | Identified in *Enterococcus faecalis* NZ_CP091229.1 (Locus tag: LQ066_14000) | Unpublished |
| Schlafen | Identified in *Enterococcus lactis* NZ_CP084245.1 (Locus tag: K9N64_RS00170) | Unpublished |
| SuperInfection Exclusion | Identified in *Enterococcus faecium* NZ_CP175931.1 (Locus tag: ACJXDE_RS13410) | Unpublished |
| Phage Head Morphogenesis | Identified in *Lactobacillus helveticus* NC_017467.1 (Locus tag: LBHH_RS08930) | Zhao *et al.,* 2011 |
| Toprim | Identified in *Enterococcus faecalis* NZ_CP091891.1 (Locus tag: L5I18_RS16595) | Unpublished |
| DUF6056 | Identified in *Enterococcus avium* NZ_CP034168.1 (EH197_RS00300) | Yu *et al.,* 2019 |
| EF_KAP D246A,D247A | Generated by ATW PCR with EF_KAP D246A, D247A F & R | This Study |
| EF_KAP Region of Unknown Function KO | Generated by two-piece ATW PCR | This Study |
| MaG KAP 1 | Identified in human fecal microbiome metagenomes | Human Microbiome Project |
| MaG KAP 2 | Identified in human fecal microbiome metagenomes | Human Microbiome Project |
| MaG KAP 3 | Identified in human fecal microbiome metagenomes | Human Microbiome Project |
| *Primers* | | |
| EF_KAP D246A, D247A F | GTATTGATCATTGCTGCTTTTGATCG | This Study |
| EF_KAP D246A, D247A R | CTTTATTTTTTAAAAAGTACGCTGAAATTG | This Study |
| EF_KAP Region of Unknown Function KO F | TAAGGATCCAGATCTTCCTTCAGGTTATG | This Study |
| EF_KAP Region of Unknown Function KO R | GTAAGCACCTTCCTGATTTAATTGAGAAATACG | This Study |
| pLZ12A non-coding region F | NNNNNNCTCGAGGTTTTTTATAAATCTATATTTAAGTAGCTTTATTGTTG | This Study |
| pLZ12A non-coding region R | NNNNNNCTCGAGGTTGGCGAAAACGTTGGCGATT | This Study |
