## Supplementary Table 3 for "Small serine recombinases are markers for antiphage defense system discovery"

**Supplemental Table 3** Domains tested in this study from the Bacillota dataset. For a full list of domains identified see Supplemental Table 2.

| Domain | InterPro ID | Adjusted P Value | N | ORF Name |
|---|---|---|---|---|
| KAP family P-loop domain | IPR011646 | 1.51E-51 | 18 | KAP NTPase / EF_KAP |
| Schlafen, AlbA_2 domain | IPR007421 | 2.05E-38 | 21 | Schlafen |
| Superinfection exclusion protein B | IPR025982 | 1.46E-16 | 4 | Superinfection Exclusion |
| Protein of unknown function DUF6056 | IPR045691 | 1.02E-4 | 3 | DUF6056 |
| Phage head morphogenesis domain | IPR006528 | 7.37E-91 | 57 | Phage Head Morphogenesis |
| TOPRIM domain | IPR006171 | 2.57E-305 | 532 | Toprim |
